# Chromatic bacteria – A broad host-range plasmid and chromosomal insertion toolbox for fluorescent protein expression in bacteria

**DOI:** 10.1101/402172

**Authors:** Rudolf O. Schlechter, Hyunwoo Jun, Michał Bernach, Simisola Oso, Erica Boyd, Dian A. Muñoz-Lintz, Renwick C. J. Dobson, Daniela M. Remus, Mitja N. P. Remus-Emsermann

**Author notes:** Both authors contributed equally to this work (Hyunwoo Jun, Michał Bernach). **Correspondence:** Mitja N. P. Remus-Emsermann.

## Abstract

Differential fluorescent labelling of bacteria has become instrumental for many aspects of microbiological research, such as the study of biofilm formation, bacterial individuality, evolution, and bacterial behaviour in complex environments. We designed a variety of plasmids, each bearing one of eight unique, constitutively expressed fluorescent protein genes in conjunction with one of four different antibiotic resistance combinations. The fluorophores mTagBFP2, mTurquoise2, sGFP2, mClover3, sYFP2, mOrange2, mScarlet-I, and mCardinal, encoding for blue, cyan, green, green-yellow, yellow, orange, red, and far-red fluorescent proteins, respectively, were combined with selectable markers conferring tetracycline, gentamicin, kanamycin, and/or chloramphenicol resistance. These constructs were cloned into three different plasmid backbones: a broad host-range plasmid, a Tn*5* transposon delivery plasmid, and a Tn*7* transposon delivery plasmid. The utility of the plasmids and transposons was tested in bacteria from the phyla Actinobacteria, Proteobacteria, and Bacteroidetes. We were able to tag representatives from the phylum Proteobacteria at least via our Tn*5* transposon delivery system. The here constructed plasmids are available to the community and provide a valuable tool to investigate bacteria-bacteria, bacteria-host, and bacteria-environmental interactions.

## Introduction

Labelling bacterial strains using fluorescent proteins have been used in manifold investigations, such as the study of biofilm formation, bacterial individuality and evolution, and bacterial behaviour in complex environments (e.g., plant surfaces, soil, or the mammalian gut) (Diard *et al.*, 2013; Kroupitski *et al.*, 2009; Monier and Lindow, 2005; Remus-Emsermann *et al.*, 2013b; Remus-Emsermann and Schlechter, 2018; Tecon and Leveau, 2012; Tolker-Nielsen and Molin, 2000; Whitaker *et al.*, 2017). Such studies require the equipment of bacteria with bright, stable, fast maturing, and highly abundant fluorescent proteins, which ideally offer unique spectral properties to unambiguously distinguish between differently labelled fluorescent bacteria. This is especially true during non-invasive *in situ* studies without additional staining procedures, such as fluorescence *in situ* hybridisation, and where autofluorescence and low signal-to-noise ratio might hinder observations (Remus-Emsermann and Schlechter, 2018).

To date, several mechanisms have been described by which fluorescent protein genes can be delivered into bacterial cells and/or integrated into their chromosomes (Andersen *et al.*, 1998; Barbier and Heath Damron, 2016; Bloemberg *et al.*, 2000; Choi and Schweizer, 2006; Lagendijk *et al.*, 2010; Lambertsen *et al.*, 2004; Ledermann *et al.*, 2015; Miller *et al.*, 2000; Remus-Emsermann *et al.*, 2016a; Schada von Borzyskowski *et al.*, 2015). Plasmids are attractive tools for fluorescent protein expression in bacteria, as they can be easily delivered into host cells and lead to high fluorescent protein production due to their presence in multiple copies (Million-Weaver *et al.*, 2012). Plasmid stability usually requires a continuous selection (e.g., antibiotics (Andersson and Hughes, 2011)) and, while only some plasmids were found to be maintained in bacterial populations without the pressure exerted by antibiotics (Bloemberg *et al.*, 2000; Rodriguez *et al.*, 2017), the absence of a selective pressure was reported to lead to plasmid loss in growing bacterial populations (Lau *et al.*, 2013; Smith and Bidochka, 1998; Summers, 1991). However, some experimental systems are not compatible with the use of antibiotics as a selective pressure. In natural environments, bacteria often occur as biofilms, where cells are fixed in space and enclosed in an extracellular polymeric substance matrix. Within a biofilm, individual cells experience heterogeneous environments, and owing to their location, they might either be exposed to or protected from antibiotic pressure (Stewart and Costerton, 2001).

To overcome the disadvantage of plasmids loss or their inability to replicate in some hosts, chromosomal insertion of fluorescent markers is advantageous. Chromosomal insertions cannot be lost in the same fashion as plasmids during division, and the probability of mutations that would disrupt the functionality of fluorescent protein genes is low. Several molecular tools and mechanisms exist to integrate chromosomal insertions into bacterial genomes, such as homologous recombination (Ledermann *et al.*, 2015), CRISPR-Cas9 (Jinek *et al.*, 2012), Zinc-finger nucleases (Carroll, 2011), or transposase-based systems (Liu *et al.*, 2013; Peters, 2014; Reznikoff, 2008).

Transposon systems can differ widely in their host specificity and mode of integration. For instance, the Tn*5* transposon has been shown to be functional in a wide range of Gram-negative bacteria and to randomly insert into their genomes with high efficiency (Reznikoff, 2008). This ability has been exploited to generate random mutations libraries (Christen *et al.*, 2011; de Lorenzo *et al.*, 1990), but also to integrate fluorescent protein genes into bacterial genomes (Andersen *et al.*, 1998; Schada von Borzyskowski *et al.*, 2015). In contrast to Tn*5* transposons, Tn*7* transposons only integrate into specific regions, such as the *att*Tn*7* site, in the host chromosome. Integration at *att*Tn*7* is mediated by four transposases, TnsA, B, C and D and appears to be prevalent in phylogenetically diverse species (Choi and Schweizer, 2006; McKenzie and Craig, 2006; Parks and Peters, 2007). Even though the host range of Tn*7* transposons is limited compared to the Tn*5* transposon, it has the tremendous advantage that the insertion does not disrupt any genes and has been argued to have little effect on bacterial fitness.

Here, we developed three sets of plasmids, which contain genes encoding for the latest generation of fluorescent proteins, i.e. that have blue, cyan, green, yellow, orange, red, and far-red fluorescence emissions, with different degrees of spectral overlap. The fluorescent protein genes mTagBFP2 (mTB2), mTurquoise2 (mTq2), sGFP2, mClover3 (mCl3), sYFP2, mOrange2 (mO2), mScarlet-I (mSc), and mCardinal (mCa) were combined with four combinations of antibiotic resistance genes, i.e. gentamicin, kanamycin, tetracycline and/or chloramphenicol, on three different plasmid backbones: a broad host-range plasmid, a Tn*5* transposon delivery plasmid, and a Tn*7* transposon delivery plasmid (Figure 1). The broad host-range plasmid contains the pBBR1 origin of replication (Miller *et al.*, 2000). The Tn*5* transposon plasmid pAG408 is based on an R6K origin of replication. It is a suicide plasmid that only replicates in presence of the π factor (Suarez *et al.*, 1997). The Tn*7* transposon plasmid pGRG36 is based on the thermo-unstable pSC101 origin of replication and is therefore a conditional suicide plasmid for species closely related to *E. coli* and a suicide plasmid in all other bacteria (McKenzie and Craig, 2006). The expression of the fluorescence protein genes is driven by the *nptII* promoter of the neomycin phosphotransferase gene (i.e. kanamycin resistance gene). The *nptII* promoter is considered constitutive, strong and was previously used to drive fluorescent protein expression from chromosomal insertions (Ledermann *et al.*, 2015; Ramirez-Mata *et al.*, 2018). We demonstrate the delivery of plasmids and transposon in a broad phylogenetic background of recently isolated environmental bacteria. Furthermore, we show the heterologous expression of fluorescent proteins at the single-cell resolution in those environmental bacterial strains using fluorescence microscopy.

**Figure 1.**
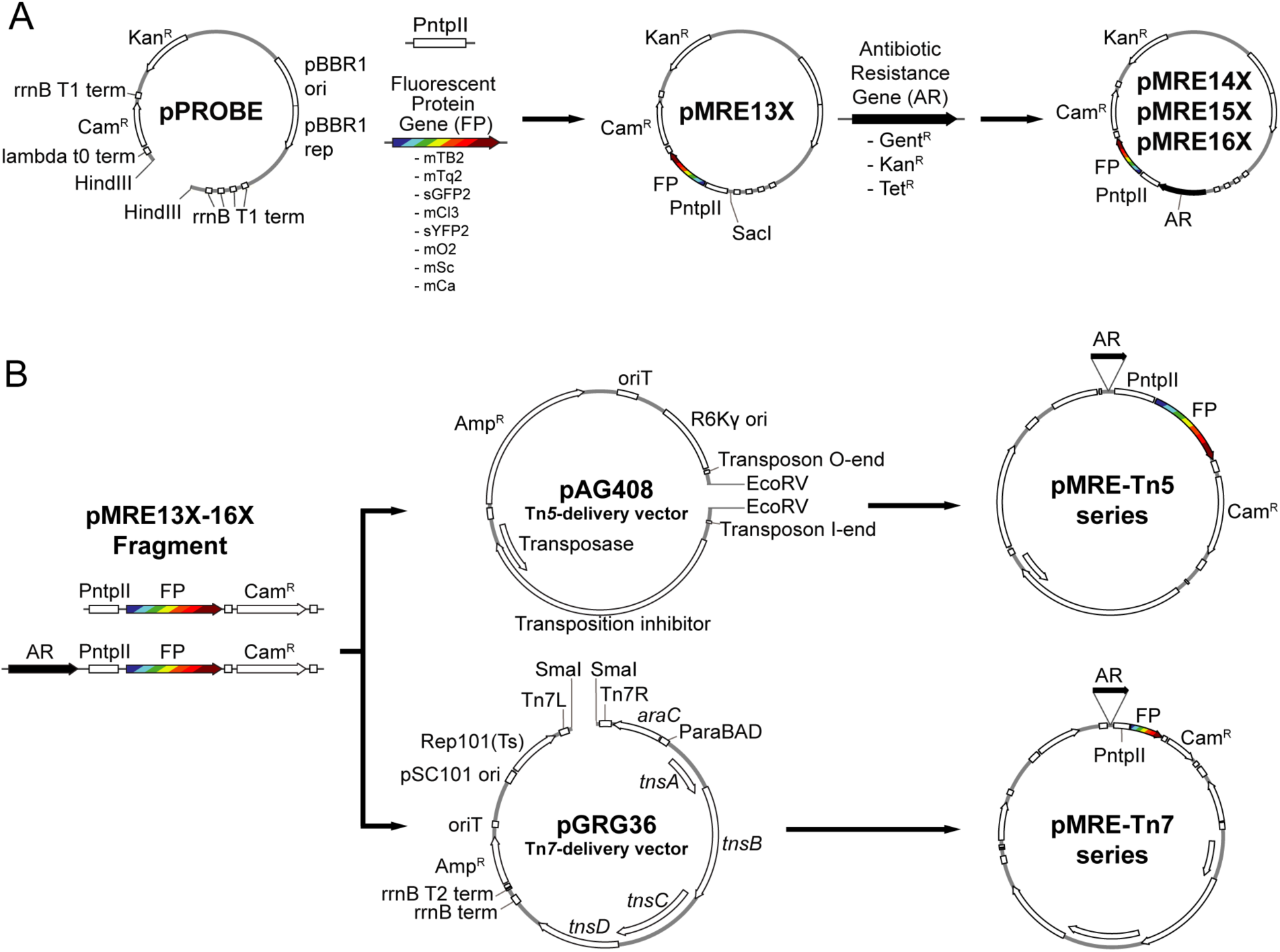
Overview of cloning procedures. (A) Construction of the pMRE plasmid series. To obtain the pMRE13X plasmid series, HindIII-digested pFru97 was isothermally assembled with a fragment containing the *ntpII* promoter (P*ntpII*) and a fragment carrying one of eight fluorescent protein genes (FP) - mTagBFP2, mTurquoise2, sGFP2, mClover3, sYFP2, mOrange2, mScarlet-I, and mCardinal. The rest of the pMRE series were constructed through the assembly of SacI-digested pMRE13X and one of three additional antibiotic resistance gene (AR) - Gentamicin (Gent^R^), Kanamycin (Kan^R^), and Tetracycline (Tet^R^). (B) Construction of the transposon delivery plasmid series pMRE-Tn5 and pMRE-Tn7. PCR products using the pMRE plasmid series as templates containing AR, FP and a Chloramphenicol resistance (Cam^R^) were isothermally inserted into EcoRV digested pAG408 to yield the pMRE-Tn5 series. Similarly, PCR products containing AR, FP and a Chloramphenicol resistance (Cam^R^) were blunt-end cloned into SmaI digested pGRG36 to yield the pMRE-Tn7 series.

## Materials and Methods

### Media and growth conditions

Bacterial strains and their growth media are shown in Table 1. Growth media were prepared according to the manufacturer’s recommendations and supplemented with 1.5 % agar (Agar No.1, Oxoid) where commercial broth media was used as a base: lysogeny broth agar (LBA, Lysogeny broth Miller, Merck), tryptic soy broth agar (TSA, Merck), nutrient agar (NA, Nutrient broth, Oxoid), or Reasoner’s 2a agar (R2A, HiMedia). *E. coli* carrying pMRE or pMRE-Tn5 plasmids were cultivated at 37 °C, whilst pMRE-Tn7 plasmid-carrying cells were cultivated at 30 °C to prevent plasmid loss. All other strains used in this study were cultivated at 30 °C. For counterselection after conjugation, auxotroph *E. coli* S17-1 was grown on either MM2 agar medium (Zengerer *et al.*, 2018) (4.0 g L^-1^ l-asparagine, 2.0 g L^-1^ K_2_HPO_4_, 0.2 g L^-1^ MgSO_4_ · 7H_2_O, 3.0 g L^-1^ NaCl, 10.0 g L^-1^ sorbitol, 15 g L^-1^ agar) or minimal agar medium (Harder *et al.*, 1973) (1.62 g L^-1^ NH_4_Cl, 0.2 g L^-1^ MgSO_4_, 1.59 g L^-1^ K_2_HPO_4_, 1.8 g L^-1^ NaH_2_PO_4_ · 2H_2_O, 15 g L^-1^ agar, with the following trace elements: 15 mg L^-1^ Na_2_EDTA_2_·H_2_O, 4.5 mg L^-1^ZnSO_4_ · 7H_2_O, 3 mg L^-1^ CoCl_2_ · 6H_2_O, 0.6 mg L^-1^ MnCl_2_, 1 mg L^-1^ H_3_BO_3_, 3.0 mg L^-1^ CaCl_2_, 0.4 mg L^-1^ Na_2_MoO_4_·2H_2_O, 3 mg L^-1^ FeSO_4_·7H_2_O, and 0.3 mg L^-1^ CuSO_4_·5H_2_O) supplemented with 0.4 % w/v succinate or glucose were used, depending on the recipient strain (Table 1). Where appropriate, media were supplemented with antibiotics in the following concentrations: 100 mg L^-1^ ampicillin, 50 mg L^-1^ kanamycin, 20 mg L^-1^ gentamicin, 15 mg L^-1^ chloramphenicol, or 15 mg L^-1^ tetracycline.

**Table 1.** Bacterial strains, their relevant naturally-occurring antibiotic resistances, standard growth media and conditions for selection after conjugation experiments.

| Strain | Relevant resistances | Growth medium | Source |
| --- | --- | --- | --- |
| *E. coli* DH5α | n.a. | LBA | Thermo Fisher Scientific |
| *E. coli* HST08 | n.a. | LBA | Stellar, Clontech |
| *E. coli* S17-1 | n.a. | LBA |  |
| *E. coli* Top10 | n.a. | LBA | Thermo Fisher Scientific |
| *Acidovorax* sp. Leaf84 | n.a. | NA/ Succinate | Bai *et al.*, 2015 |
| *Bradyrhizobium* sp. Leaf396 | n.a. | R2A/ Succinate | Bai *et al.*, 2015) |
| *Erwinia amylovora* CFBP1430S | n.a. | TSA/ Succinate | Smits *et al.*, 2010 |
| *Methylobacterium* sp. Leaf92 | n.a. | R2A/ Succinate | Bai *et al.*, 2015 |
| *Microbacterium* sp. Leaf320 | Gent<sup>R</sup> | NA/ Glucose | Bai *et al.*, 2015 |
| *Pantoea agglomerans* 299R | Cam<sup>R</sup> | TSA/ Succinate | Remus-Emsermann *et al.*, 2013a |
| *Pedobacter* sp. Leaf194 | Gent<sup>R</sup>, Kan<sup>R</sup> | R2A/ R2A + Gent | Bai *et al.*, 2015 |
| *Pseudomonas syringae* B728a | Cam<sup>R</sup> | TSA/ Succinate | Feil *et al.*, 2005 |
| *Pseudomonas citronellolis* P3B5 | Cam<sup>R</sup> | NA/ Succinate | Remus-Emsermann *et al.*, 2016b |
| *Sphingomonas melonis* FR1 | n.a. | NA/ Succinate | Innerebner *et al.*, 2011 |
| *Rhodococcus* sp. Leaf225 | n.a. | NA/ Glucose | Bai *et al.*, 2015 |

n.a.: not applicable

### Plasmid construction

All plasmids used or constructed in this study are shown in Table 2. Generic plasmid maps and cloning procedures are presented in Figure 1. For plasmid construction, PCRs were performed using Phusion High-Fidelity DNA polymerase (Thermo Scientific) following the manufacturer’s recommendations. Annealing temperatures (Ta) were chosen based on the respective melting temperature (*t*_m_) of the primers (Table 3). Touchdown PCRs were performed to amplify PCR products with overlapping ends for isothermal assemblies. To that end, the initial Ta was set to 10 °C above the lowest *t*_m_ of the respective primers and the Ta was gradually reduced 1 °C per cycle for a total of ten cycles. After the tenth cycle, a 2-step PCR with Ta set to 72 °C was performed for 25 cycles.

**Table 2.** Plasmids used in this work.

| Name | Notable features | Source |
| --- | --- | --- |
| pFru97 | Kan<sup>R</sup>, Cam<sup>R</sup> | Tecon and Leveau, 2012 |
| pGRG36 | Tn7 transposon, Amp<sup>R</sup> | McKenzie and Craig, 2006 |
| pAG408 | Tn5 transposon, Amp<sup>R</sup>, Gent<sup>R</sup>, Kan<sup>R</sup> | Suarez *et al.*, 1997 |
| mTagBFP2-pBAD | mTagBFP2 | Subach *et al.*, 2011 |
| pLifeAct-mTurquoise2 | mTurquoise2 | Goedhart *et al.*, 2012 |
| pMREtn5-Ptuf-sYFP2 | sYFP2 | This work |
| mOrange2-pBAD | mOrange2 | Michael Davidson lab |
| pTriEx-RhoA-wt_mScarlet-I_SGFP2 | mScarlet-I, sGFP2 | Bindels *et al.*, 2017 |
| mCardinal-pBAD | mCardinal | Chu *et al.*, 2014 |
| pRJpahp-gfp+ | nptII promoter | Ledermann *et al.*, 2015 |
| pTE105-mChe | Tet<sup>R</sup> | Schada von Borzyskowski *et al.*, 2015 |
| pMRE13X | Kan<sup>R</sup>, Cam<sup>R</sup> | This work |
| pMRE14X | Kan<sup>R</sup>, Cam<sup>R</sup>, Gent<sup>R</sup> | This work |
| pMRE15X | Kan<sup>R</sup>, Cam<sup>R</sup>, Kan<sup>R</sup> | This work |
| pMRE16X | Kan<sup>R</sup>, Cam<sup>R</sup>, Tet<sup>R</sup> | This work |
| pMRE-Tn7-13X | Tn7 transposon, Amp<sup>R</sup>, Cam<sup>R</sup> | This work |
| pMRE-Tn7-14X | Tn7 transposon, Amp<sup>R</sup>, Cam<sup>R</sup>, Gent<sup>R</sup> | This work |
| pMRE-Tn7-15X | Tn7 transposon, Amp<sup>R</sup>, Cam<sup>R</sup>, Kan<sup>R</sup> | This work |
| pMRE-Tn7-16X | Tn7 transposon, Amp<sup>R</sup>, Cam<sup>R</sup>, Tet<sup>R</sup> | This work |
| pMRE-Tn5-13X | Tn5 transposon, Amp<sup>R</sup>, Cam<sup>R</sup> | This work |
| pMRE-Tn5-14X | Tn5 transposon, Amp<sup>R</sup>, Cam<sup>R</sup>, Gent<sup>R</sup> | This work |
| pMRE-Tn5-15X | Tn5 transposon, Amp<sup>R</sup>, Cam<sup>R</sup>, Kan<sup>R</sup> | This work |
| pMRE-Tn5-16X | Tn5 transposon, Amp<sup>R</sup>, Cam<sup>R</sup>, Tet<sup>R</sup> | This work |

X is a placeholder for either 0, 1, 2, 3, 4, 5, 6, or 7 representing mTagBFP2, mTurquoise, sGFP2, sYFP2, mOrange, mScarlet-I, mCardinal, or mClover3 encoding versions of the plasmids, respectively.

**Table 3.**
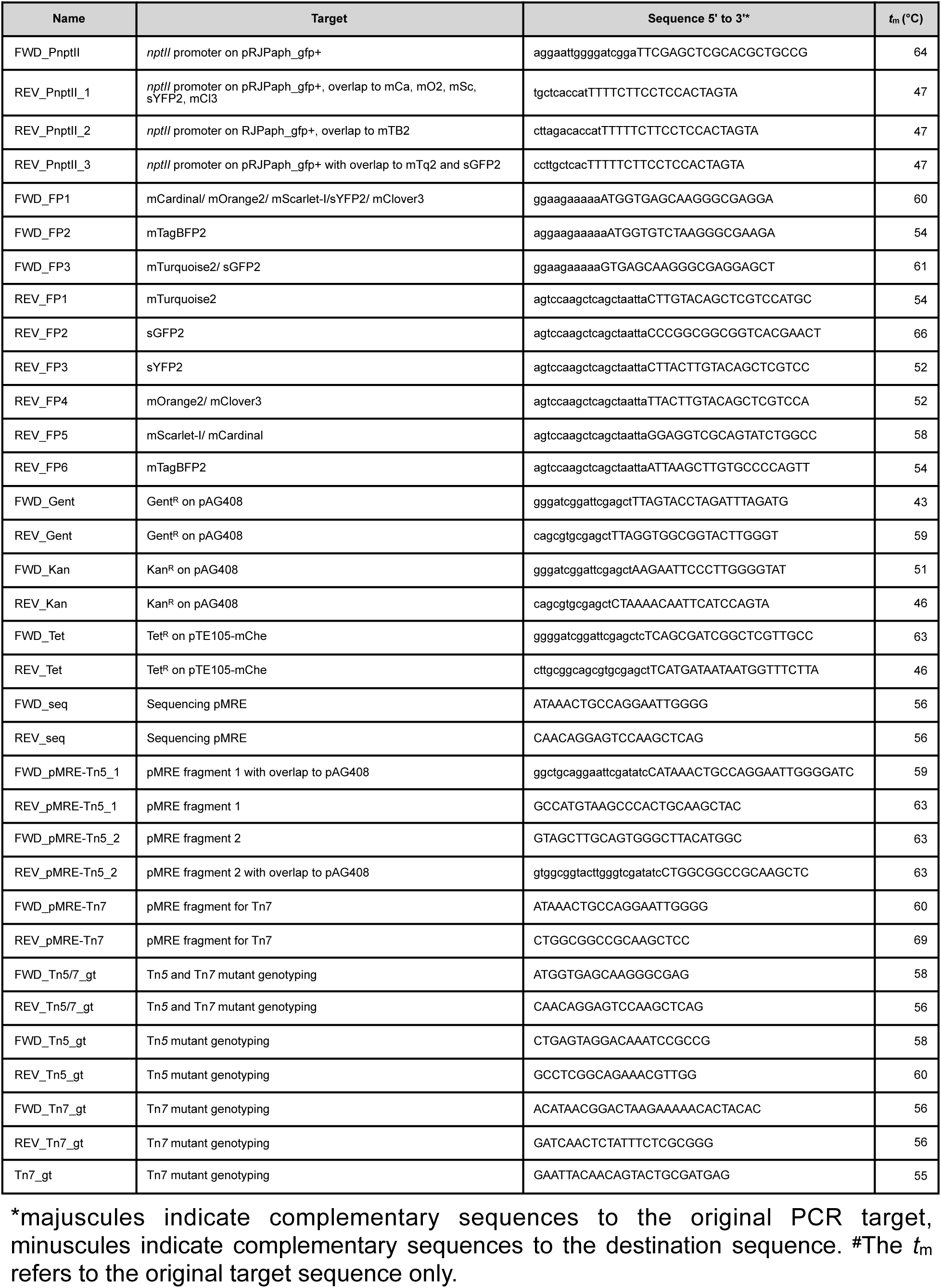
Primers used in this work.

Several plasmids that were used as source for fluorescent protein genes were acquired from the plasmid repository addgene.org: pGRG36 was a gift from Nancy Craig (Addgene plasmid #16666). pTriEx-RhoA-wt_mScarlet-I_SGFP2 (Addgene plasmid #85071) and pLifeAct-mTurquoise2 (Addgene plasmid #36201) were gifts from Dorus Gadella. pNCS-mClover3 was a gift from Michael Lin (Addgene plasmid #74236). Plasmids mCardinal-pBAD (Addgene plasmid #54800) and mTagBFP2-pBAD (Addgene plasmid #54572) were gifts from Michael Davidson and mOrange2-pBAD was a gift from Michael Davidson & Roger Tsien (Addgene plasmid #54531). pFru97 was a gift from Johan Leveau (Tecon and Leveau, 2012). All plasmids that were used for PCR amplification were isolated using the DNA-spin plasmid DNA purification kit (INtRON biotechnology) following the manufacturer’s recommendations. The designated plasmid backbones of the herein constructed plasmids, pFru97, pGRG36, and pAG408, were isolated using the Zyppy™ plasmid midiprep kit (Zymo) following the manufacturer’s recommendations. Gel purification and PCR clean-up was performed using the Monarch DNA Gel Extraction Kit (NEB) or the DNA Clean & Concentrator™-5 Kit (Zymo), respectively.

To construct the pMRE130 series, the promoter of the *nptII* gene (PnptII) was amplified from pRJPaph_gfp+ using primer FWD_PnptII, which contains overlapping sequences to HindIII-digested pFru97, and either primer REV_PnptII_1, REV_PnptII_2, or REV_PnptII_3, which contain overlapping sequences to the different fluorescent protein genes. The fluorescent protein genes were amplified using primer FWD_FP1, FWD_FP2, or FWD_FP3 with overlapping sequences to the PnptII fragment and primer X is a placeholder for either 0, 1, 2, 3, 4, 5, 6, or 7 representing mTagBFP2, mTurquoise, sGFP2, sYFP2, mOrange, mScarlet-I, mCardinal, or mClover3 encoding versions of the plasmids, respectively.

REV_FP1, REV_FP2, REV_FP3, REV_FP4, REV_FP5, or REV_FP6 with overlapping sequences to HindIII-digested pFru97. The two resulting fragments were mixed with the HindIII-digested pFru97 backbone and isothermally assembled to yield plasmids pMRE130, pMRE131, pMRE132, pMRE133, pMRE134, pMRE135, pMRE136, and pMRE137. To construct the pMRE140 plasmid series, the gentamicin resistance gene (Gent^R^) was amplified from pAG408 using primers FWD_Gent and REV_Gent with overlaps to SacI-digested plasmids of the p MRE 13 X series, yielding pMRE140,pMRE141, pMRE142, pMRE143, pMRE144, pMRE145, pMRE146, and pMRE147. To construct the pMRE150 plasmid series, the kanamycin resistance gene (Kan^R^) was amplified from pAG408 using primers FWD_Kan and REV_Kan with overlapping sequences to SacI-digested plasmids of the pMRE13X series, yielding pMRE150, pMRE151, pMRE152, pMRE153, pMRE154, pMRE155, pMRE156, and pMRE157. To construct the pMRE16X plasmid series, the tetracycline resistance genes (Tet^R^) were amplified from pTE105-mChe using primers FWD_Tet and REV_Tet with overlapping sequences to SacI-digested plasmids of the pMRE13X series, yielding pMRE160, pMRE161, pMRE162, pMRE163, pMRE164, pMRE165, pMRE166, and pMRE167.

Isothermal assembly was performed as described previously (Benoit *et al.*, 2016; Gibson *et al.*, 2009; Remus-Emsermann *et al.*, 2016a). In short, plasmid backbones and inserts with overlapping sequences were mixed in a 1:3 molar ratio (20-100 ng backbone) to reach a total volume of 5 µL. Then, 15 µL of isothermal assembly mix were added and the reaction was incubated for 15 minutes at 50 °C. Transformation of chemically competent *E. coli* strains was performed using standard procedures as recommended by the respective suppliers or as described in Sambrook *et al.* (Sambrook *et al.*, 1989) (See Table 1).

To construct the pMRE-Tn5 plasmid series, pMRE1XX plasmids were used as a template to amplify the desired DNA fragments, which include one of the eight fluorescent protein gene and one of the four antibiotic resistance marker combinations. Cassettes were amplified as two fragments using the primers FWD1_pMRE_pAG408 and RV1_pMRE_pAG408 or FWD2_pMRE_pAG408 and RV2_pMRE_pAG408. Primers FWD1_pMRE_pAG408 and RV2_pMRE_pAG408 contain overlapping sequences to EcoRV-digested pAG408. For isothermal assemblies, EcoRV-digested pAG408 was mixed with the insert fragments in a 1:3:3 ratio as described above and assembled plasmids were transformed into chemically competent *E. coli* S17-1.

To construct the pMRE-Tn7 family, fluorescent protein genes and antibiotic resistance genes were amplified from the pMRE1XX series using primers FWD_pMRE-Tn7 and REV_pMRE-Tn7. The amplicons were gel purified using the Monarch DNA Gel Extraction Kit (NEB) and phosphorylated using T4 polynucleotide kinase (Life Technologies). pGRG36 was SmaI digested and dephosphorylated using thermosensitive alkaline phosphatase according to the manufacturers’ recommendations (Fast AP, Life Technologies). Following dephosphorylation, plasmids were purified using the DNA Clean & Concentrator™-5 Kit (Zymo). Amplicons were then cloned into linearised and dephosphorylated pGRG36 using Quick-Stick T4 Ligase (Bioline). Ligations were purified using the DNA Clean & Concentrator™-5 Kit and transformed into chemically competent *E. coli* (Top10, One Shot™ MAX Efficiency™ DH5α™-T1R, or Stellar).

All pMRE1XX series plasmids were verified by Sanger sequencing using primers FWD_seq and REV_seq at Macrogen. pMRE-Tn5-1XX and pMRE-Tn7-1XX series plasmids were verified by PvuII restriction digests at 37 °C for 1 h. To provide a convenient means of plasmid delivery, all plasmids were subsequently transformed into *E. coli* S17-1, which allows conjugations to recipient strains.

### Conjugation

Recipients were grown on standard agar media for up to four days, depending on the growth rate of each environmental strain (see Table 1). Donor strains *E. coli* S17-1 were grown on LBA overnight (see Table 1). Plasmids delivered into recipient strains include pMRE135, pMRE145, pMRE165, pMRE-Tn5-143, pMRE-Tn5-145, pMRE-Tn5-165, pMRE-Tn7-145, or pMRE-Tn7-165. Freshly grown bacteria were harvested using a loop and resuspended in 1 × phosphate buffered saline (1 × PBS (8 g L^-1^ NaCl, 0.24 g L^-1^ KCl, 1.42 g L^-1^ Na_2_HPO_4_, 0.24 g L^-1^ KH_2_PO_4_)) to reach an OD_600nm_ of 1. Each recipient strain was mixed with their respective donor strains in 1:1 ratio and the mix was then concentrated by centrifugation (4000 *g*, 5 min) to reach an estimated OD_600nm_ of 20. Bacterial mixes were drop spotted on NA and incubated for 18 hours at 30 °C. For conjugations using pMRE-Tn7, 0.1 % w/v arabinose was added to the medium, as the Tn*7* transposase genes are under the control of an arabinose-inducible promoter. After incubation, the cells were harvested and resuspended in 1 mL 1 × PBS. The bacterial mix was spread onto appropriate minimal agar media containing a sole carbon source (either 0.4 % w/v succinate or 0.4 % w/v glucose, see Table 1) and appropriate antibiotics. Depending on the recipient strain, transconjugants appeared within 5-10 days. To further counterselect against *E. coli*, single colonies were restreaked at least twice onto fresh minimal media before growing them on complex media.

### Validation of transposon insertion events

To provide a convenient tool to assess successful insertions of Tn*5* and Tn*7* transposons, a multiplex PCR was designed to determine the presence of plasmid backbones and fluorescent protein genes in one reaction. Primers FWD_Tn5/7_gt and RV_Tn5/7_gt were designed to amplify the fluorescent protein genes for both Tn*5* and Tn*7* delivery systems, while the combinations FWD_Tn5_gt and RV_Tn5_gt and FWD_Tn7_gt, RV_Tn7_gt, and Tn7_gt amplify a specific fragment of the backbone of Tn*5* and Tn*7* plasmids, respectively. For Tn*5* insertions, the primer mix FWD_tn5/7_gt, RV_tn5/7_gt, FWD_tn5_gt, and RV_tn5_gt was used. For Tn*7* insertions, primer mix FWD_tn5/7_gt and RV_tn5/7_gt and primer mix FWD_tn7_gt, RV_tn7_gt, and Tn7_gt were used. Reactions were performed using the KAPA2G Fast HotStart ReadyMix PCR kit (Kapa Biosystems) following the manufacturer’s instructions.

### Fluorescent protein absorption and emission spectra and fluorescence intensity

The absorption and emission spectra of the different proteins were measured in a Cary Eclipse Fluorescence Spectrophotometer (Agilent Technologies). To that end, overnight cultures of *E. coli* strains expressing the different fluorescent proteins were harvested by centrifugation (4000 *g*, 3 min), washed in 1 × PBS and resuspended to reach an OD_600nm_ of 1. Then, 1 mL of each culture was measured in a polystyrene cuvette (VWR), besides cultures expressing mTagBFP2 and mCardinal, which were measured in a far-UV quartz cuvette (Agilent Technologies) due to their spectral properties. Absorption and emission were measured in the spectrophotometer in 1 nm intervals. The absorption and emission spectra of the proteins used in this study are available as “protein collection” on the webpage fpbase.org (https://www.fpbase.org/collection/78/; (Lambert, 2018)). This online platform allows for a convenient comparison of existing microscopy hardware and fluorophore properties.

To determine brightness of *E. coli* DH5α carrying plasmids or transposon insertions, respective strains were grown in eight biological replicates in LB broth (300 rpm, 37 °C) to reach an OD_600nm_ of 1. Fluorescence intensity of cultures were measured in a flat bottom 96-well plate (Costar) using a FLUOstar Omega plate reader (BMG Labtech) equipped with an excitation filter (band pass 580/10) and an emission filter (band pass 620/10). Fluorescence intensity was corrected by background subtraction.

### Fluorescence microscopy and image processing

Fluorescent protein-expressing *E. coli* were grown overnight in LB broth (210 rpm, 37 °C). Cultures were harvested by centrifugation (4000 *g*, 3 min) and washed in 1 × PBS. Bacterial cultures were resuspended in 1 × PBS to an OD_600nm_ of 0.01, i.e. ∼10^7^ bacteria/mL, and were mounted on agarose slabs (0.8 % w/v agarose in H_2_O). Fluorescence microscopy was performed on a Zeiss AxioImager.M1 fluorescent widefield microscope equipped with Zeiss filter sets 38HE, 43HE, 46HE, 47HE, and 49 (BP 470/40-FT 495-BP 525/50, BP 550/25-FT 570-BP 605/70, BP 500/25-FT 515-BP 535/30, BP 436/25-FT 455-BP 480/40, and G 365-FT 395-BP 445/50, respectively), an Axiocam 506 and the software Zeiss Zen 2.3. For confocal microscopy, a Leica SP5 confocal laser scanning microscope equipped with a 405 nm UV laser, an Argon laser line 458, 476, 514, nm, and a diode-pumped solid-state laser line 561 nm and software Leica LAS AF (version 2.6.3.8173) was used. Suspensions containing six or seven different fluorescent protein-tagged *E. coli* were detected through confocal microscopy using sequential scanning mode. For six colour mixtures consisting of mTB2, mTq2, mCl3, mO2, mSc, and mCa-expressing *E. coli*, a first scan using laser line 561 nm was used for emission windows of 570-599 nm, 600-630 nm, and 650-750 nm. Then, a second scan using laser lines 405, 458 and 476 nm were used for emission windows 440-460 nm, 480-500 nm, and 520-540 nm. For seven colour mixtures consisting of mTB2, mTq2, sGFP2, sYFP2, mO2, mSc, and mCa-expressing *E. coli*, a first scan using laser line 561 nm was used for emission windows of 570-599 nm, 600-630 nm, and 650-750 nm. A second scan using line 514 nm was used for the emission windows 540-560 nm. A third scan using lines 405, 458, 476 nm, and were used for the emission windows 440-460 nm, 480-500 nm, and 520-540 nm.

All image analysis and processing were carried out in ImageJ/FIJI (Schindelin *et al.*, 2012). Signal bleed-through was corrected using a channel subtraction method. First, individual channels were thresholded (ImageJ/Fiji threshold using Otsu) to produce binary images. Binary template images were then dilated (ImageJ/Fiji command dilate) to accommodate spherical aberrations and subtracted from channels with signal bleed-through. All channels were background subtracted.

Single-cell fluorescence was analysed as described previously (Remus-Emsermann *et al.*, 2016a). In short, bacteria were mounted on an agar slab as described above and samples were analysed using a Zeiss AxioImager.M1 at 100x magnification. Multichannel images were acquired using phase contrast and appropriate fluorescent filters (see above). Using ImageJ/Fiji, phase contrast channels were thresholded using standard settings and used as a mask to determine the mean fluorescence signal of individual particles in the respective fluorescent channels.

## Results

In this work, a total of 96 plasmids were constructed (Table 2). The plasmids constitute three sets, including *i*) 32 broad-host range plasmids based on the pBBR1 origin of replication, *ii*) 32 suicide plasmids based on the R6K origin of replication, which are maintained solely in hosts carrying the π factor, and a Tn*5*-based transposon system, and *iii*) 32 narrow host-range plasmids harbouring a pSC101 temperature-sensitive origin of replication and the Tn*7*-based transposon system. Each set consists of plasmids that carry one of eight different fluorescent protein genes and one of four different combinations of antibiotic resistances. We used a simple naming convention and separated the plasmid into the pMRE13X-series, which all confer chloramphenicol resistance, the pMRE14X-series, which all confer gentamicin and chloramphenicol resistance, the pMRE15X-series, which all confer kanamycin and chloramphenicol resistance, and the pMRE16X-series, which all confer tetracycline and chloramphenicol resistance. The pFru97-based pMRE series contains an additional kanamycin resistance. Furthermore, we named plasmids based on the contained fluorescent protein, i.e. pMRE1X0 denotes mTagBFP2, pMRE1X1 denotes mTurquoise2, pMRE1X2 denotes sGFP2, pMRE1X3 denotes sYFP2, pMRE1X4 denotes mOrange2, pMRE1X5 denotes mScarlet-I, pMRE1X6 denotes mCardinal, pMRE1X7 denotes mClover3. Transposons are indicated as pMRE-Tn5-1XX for Tn*5* transposons and pMRE-Tn7-1XX for Tn*7* transposons.

Fluorescent protein-expressing bacteria were found to exhibit the expected fluorescence absorption and emission spectra (Figure 2) (Bindels *et al.*, 2017; Chu *et al.*, 2014; Goedhart *et al.*, 2012; Kremers *et al.*, 2007; Subach *et al.*, 2011), suggesting that no changes in their spectral properties were introducing by cloning procedures. However, it was noticeable that bacterial cells harbouring the pMRE14X-series exhibited increased fluorescence intensities than in the rest of the plasmid series, likely due to readthrough of the gentamicin promoter which was cloned upstream the fluorescent protein gene. Therefore, the fluorescence intensity of the pMRE-series carrying the mScarlet-I gene in combination with all antibiotic resistance genes was determined in *E. coli* DH5α. Additionally, the emitted fluorescence was assessed in four independent *E. coli* DH5α::Tn5-MRE145 Tn*5* insertions mutants and four independent *E. coli* DH5α::Tn7-MRE145 Tn*7* insertion mutants. Plasmids pMRE135, pMRE155 and pMRE165 led to similar fluorescence in *E. coli* DH5α, however, plasmid pMRE145 resulted in significantly brighter fluorescence intensities (Figure 3A). After Tn*5* and Tn*7* transposition, *E. coli* DH5α exhibited similar fluorescence intensities, however, transposon-conferred fluorescence intensities were significantly lower than plasmid-conferred fluorescence intensities (Figure 3B).

**Figure 2.**
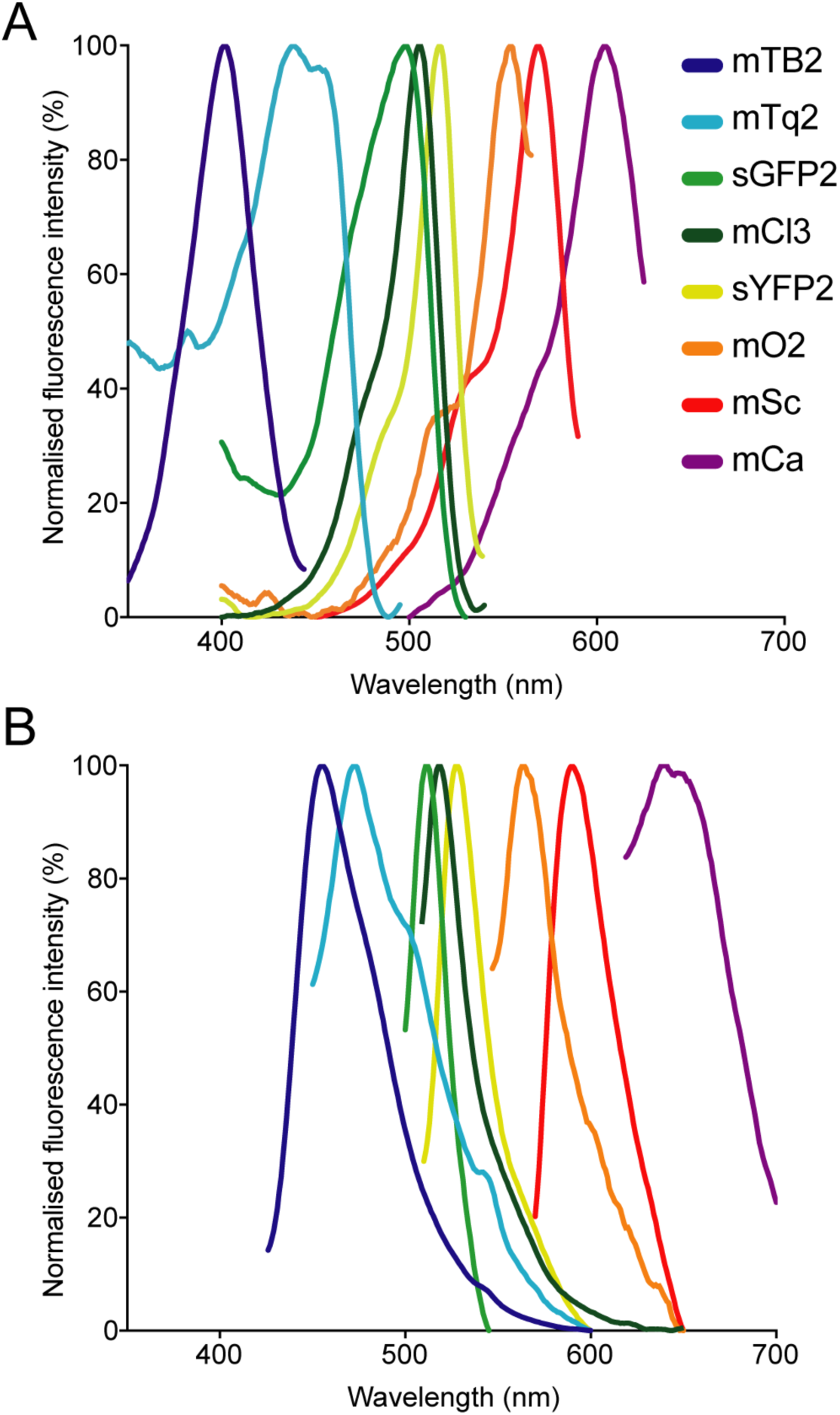
Normalised (A) absorption and (B) emission spectra of the fluorescent proteins used in this work.

**Figure 3.**
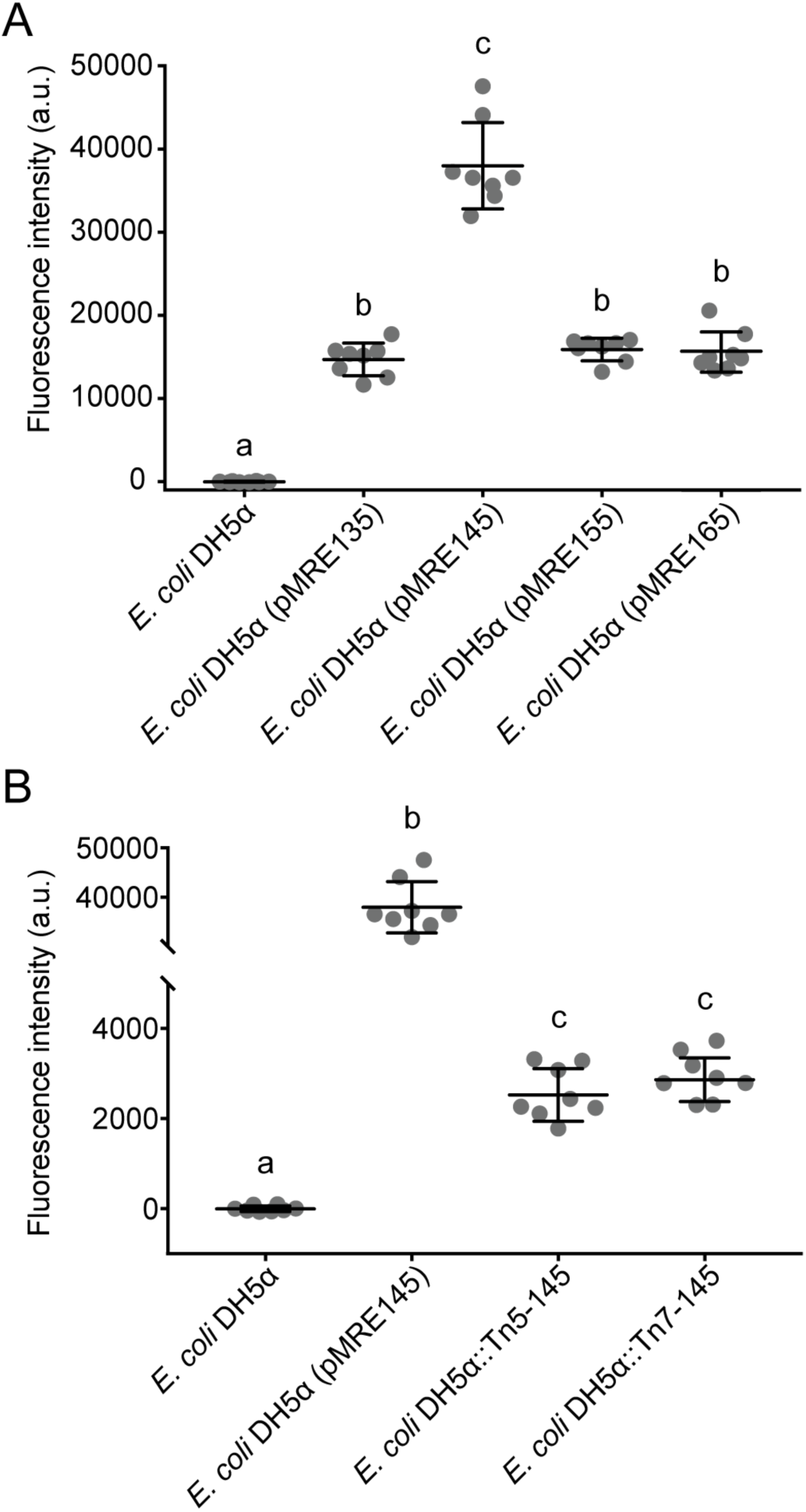
(A) Fluorescence intensity of *E. coli* DH5α cultures expressing mScarlet-I from different pMRE plasmid series (pMRE135, pMRE145, pMRE155, pMRE165). (B) Fluorescence intensity of mScarlet-I in plasmid-borne *E. coli* DH5α cultures (pMRE145), Tn*5-* and Tn*7*-insertion mutants. *E. coli* DH5α was used as control. One-way ANOVA and Tukey’s *post hoc* test were used to infer statistical differences between groups. Letters indicate differences among means with a 99.9% confidence interval (*p* < 0.0001) A.u.: arbitrary units.

### Plasmid sets are broadly transmissible into a variety of Gram-negative bacteria

Plasmid functionality and fluorescence protein expression was tested in a phylogenetically broad range of bacteria that were recently isolated from plant leaves (Table 4). The pMRE-series plasmids were successfully conjugated and maintained in *Sphingomonas melonis* FR1, *Erwinia amylovora* CFBP1430S, *Pantoea agglomerans* 299R, and *P. syringae* B728a. Tn*5* transposon plasmid were delivered by conjugation and integrated into the genome of *Bradyrhizobium* sp. Leaf396, *Methylobacterium* sp.

**Table 4.** Bacterial strains used as recipients for conjugation of pMRE, pMRE-Tn5 and pMRE-Tn7.

| Class | Order | Family | Genus species | pMRE | ::MRE-Tn5 | ::MRE-Tn7 |
| --- | --- | --- | --- | --- | --- | --- |
| Actinobacteria | Actinomycetales | Microbacteriaceae | *Microbacterium* sp. Leaf320 | no | no | no |
| Actinobacteria | Actinomycetales | Nocardiaceae | *Rhodococcus* sp. Leaf225 | no | no | no |
| α-Proteobacteria | Rhizobiales | Rhizobiaceae | *Bradyrhizobium* sp. Leaf396 | no | 165 | no |
| α-Proteobacteria | Rhizobiales | Rhizobiaceae | *Methylobacterium* sp. Leaf92 | no | 165 | no |
| α-Proteobacteria | Sphingomonadales | Sphingomonadaceae | *Sphingomonas melonis* Fr1 | 145 | 145 | 145 |
| α-Proteobacteria | Sphingomonadales | Sphingomonadaceae | *Sphingomonas phyllosphaerae* FA2 | 135 | 145 | no |
| β-Proteobacteria | Burkholderiales | Comamonadaceae | *Acidovorax* sp. Leaf84 | no | no | no |
| γ-Proteobacteria | Enterobacteriales | Enterobacteriaceae | *Erwinia amylovora* CFBP1430S | 135 | 145 | 145 |
| γ-Proteobacteria | Enterobacteriales | Enterobacteriaceae | *Pantoea agglomerans* 299R | 135 | 145 | 145 |
| γ-Proteobacteria | Pseudomonadales | Pseudomonadaceae | *Pseudomonas citronellolis* P3B5 | 145 | 145 | no |
| γ-Proteobacteria | Pseudomonadales | Pseudomonadaceae | *Pseudomonas syringae* B728a | 145 | 145 | no |
| Sphingobacteria | Sphingobacteriales | Sphingobacteriaceae | *Pedobacter* sp. Leaf194 | no | no | no |

The specific plasmids or transposons used for each strain are depicted by their ending (refer to Table 2).

Leaf92, *S. melonis* FR1, *Sphingomonas phyllosphaerae* FA2, *E. amylovora* CFBP1430S, *P. agglomerans* 299R, and *Pseudomonas citronellolis* P3B5. Tn*7* transposon plasmids were delivered by conjugation and integrated into the genome of *S. melonis* FR1, *E. amylovora* CFBP1430S, and *P. agglomerans* 299R.

For a fast assessment of transposon-mediated insertions, multiplex PCRs were developed to amplify a region of the insert (i.e. fluorescent protein gene) and plasmid backbone simultaneously, i.e. a PCR on the donor plasmid resulted in two PCR products, while a genomic integration of the transposon resulted in only one PCR product. As expected, successfully integrated insertions into the genome of all tested strains were corroborated, as no plasmid backbone was detected whilst the fluorescent protein gene was correctly amplified (Supplementary Figure 1).

### The provided fluorescent protein toolbox allows to track multiple bacterial strains in parallel

Using standard fluorescence microscopy equipment, it was possible to track up to seven bacterial strains in parallel. Employing widefield fluorescence microscopy and widely distributed fluorescence filter sets (i.e. standard blue, green and red fluorescence emission filters) three bacterial strains could be unambiguously identified. Thereby allowing to monitor three strains equipped with mTB2, mCl3, and mSc (Figure 4 A), respectively. Less common combination of filters (i.e. blue, cyan, yellow, and red fluorescence emission filters) allows monitoring of up to four strains expressing mTB2, mTq2, sYFP2, and mSc, respectively (Figure 4 B).

**Figure 4.**
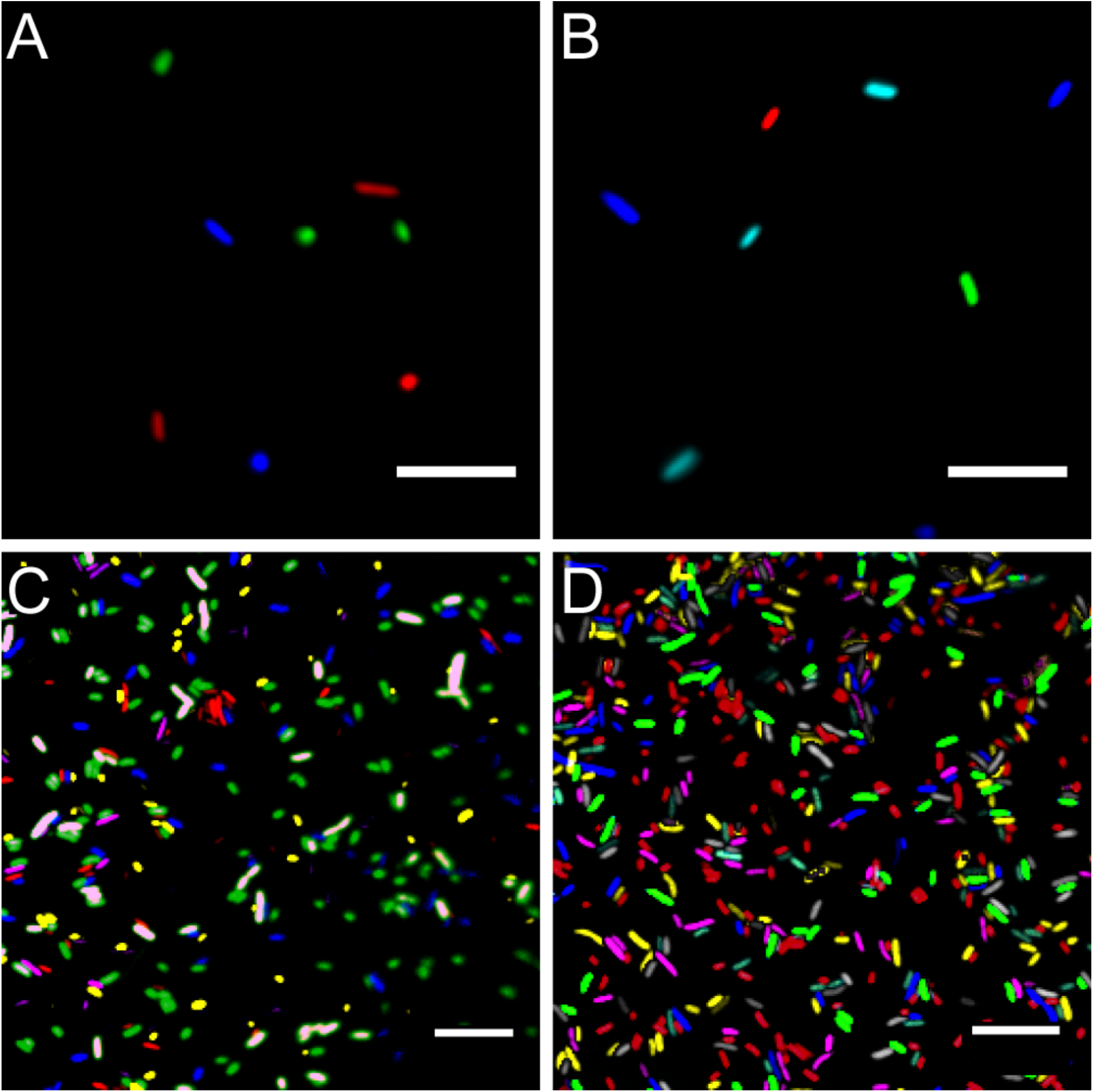
Microscopy images of fluorescent bacteria. (A) widefield epifluorescence micrograph of a *E. coli* expressing either mTB2 (blue), mCl3 (green), or mSc (red). (B) widefield epifluorescence micrograph of a *E. coli* expressing either mTB2 (blue), mTq2 (cyan), sYFP2 (green), or mSc (red). (C) Confocal microscopy of a mixed *E. coli* expressing either mTB2 (blue), mTq2 (magenta), mCl3 (yellow), mO2 (green), mSc (white), or mCa (red). (D) Confocal microscopy of a mixed *E. coli* expressing either mTB2 (cyan), mTq2 (magenta), sGFP2 (yellow), sYFP2 (gray), mO2 (red), mSc (green), or mCa (blue). In all cases, *E. coli* harboring pMRE14X plasmids series were used. Scale bar represents 10 µm.

Using filter free confocal laser scanning microscopy, it was possible to detect six different fluorescently tagged cell populations without further image processing (Figure 4 C). After bleedthrough correction (see Material and Methods, i.e. for sYFP2 and sGFP2, mO2 and mSc, and mSc and mCa) seven different fluorescently tagged cell populations could be distinguished (Figure 4D).

### A broad host range of environmental bacteria express fluorescent proteins from plasmids and transposon insertions

All bacteria that were successfully equipped with plasmids or transposons (Table 4) were grown on agar media plates and epifluorescence widefield microscopy was performed to determine fluorescence at the single-cell resolution. All bacterial strains exhibited fluorescence that could be determined at the single-cell resolution with a high signal-to-noise ratio (Figure 5). The range of fluorescence intensity of the environmental strains was comparable to *E. coli* DH5α and yielded signals in the same magnitude. Of the investigated bacteria strains, *P. syringae* B728a yielded an exceptionally bright signal (Supplementary Figure 2).

**Figure 5.**
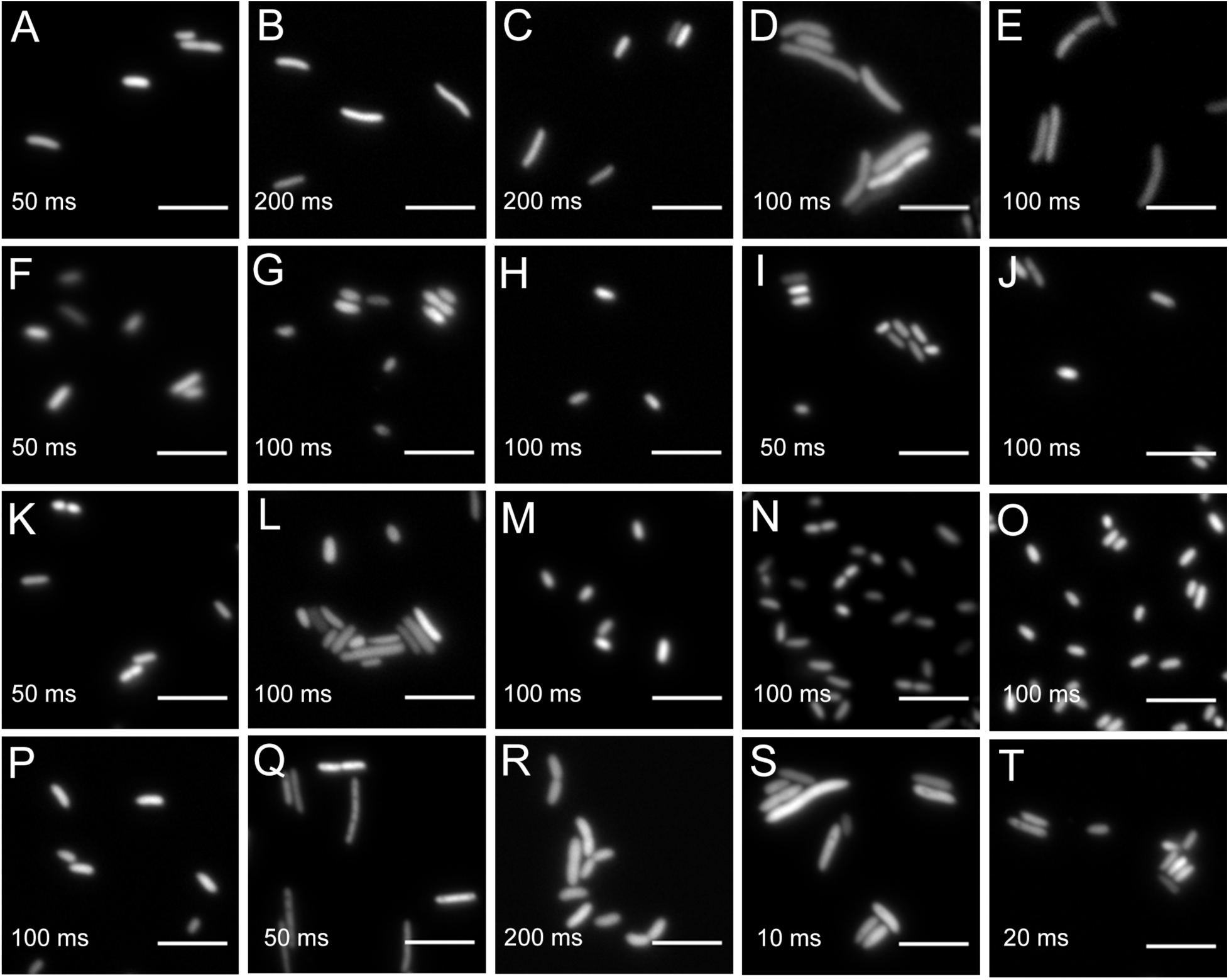
Widefield microscopy of environmental bacteria expressing fluorescent proteins. (A) *Escherichia coli* DH5α (pMRE145). (B) *E. coli* DH5α::MRE-Tn5-145. (C) *E. coli* DH5α::MRE-Tn7-145. (D) *Bradyrhizobium* sp. Leaf396::MRE-Tn5-165. (E) *Methylobacterium* sp. Leaf92::MRE-Tn5-165. (F) *Sphingomonas melonis* FR1 (pMRE145). (G) *S. melonis* FR1::MRE-Tn5-145. (H) *S. melonis* FR1::MRE-Tn7-145. (I) *Sphingomonas phyllosphaerae* FA2 (pMRE135). (J) *S. phyllosphaerae* FA2::MRE-Tn5-145. (K) *Erwinia amylovora* CFBP1430S (pMRE135). (L) *E. amylovora* CFBP1430S::MRE-Tn5-145. (M) *E. amylovora* CFBP1430S::MRE-Tn7-145. (N) *Pantoea agglomerans* 299R (pMRE135). (O) *P. agglomerans* 299R::MRE-Tn5-145. (P) *P. agglomerans* 299R::MRE-Tn7-145. (Q) *Pseudomonas citronellolis* P3B5 (pMRE145). (R*) P. citronellolis* P3B5::MRE-Tn5-145. (S) *Pseudomonas syringae* pv. *syringae* B728a (pMRE145). (T) *P. syringae* B728a::MRE-Tn5-145. Exposure times used during image acquisition are depicted in the corresponding images. Scale bars represent 5 µm.

## Discussion

A set of plasmids was constructed conferring eight fluorescent phenotypes in a wide range of bacteria. The fluorescent proteins used in this work were selected to have minimal spectral overlap, which allows parallel detection of up to four fluorescent proteins on a standard widefield microscopy system and seven fluorescent proteins on filter free confocal microscopy systems. Additionally, the series of plasmids and transposons carry four different antibiotic resistance combinations which allows the selection of bacterial mutants while accommodating their requirements of naturally-occurring antibiotic resistance.

The strong, constitutive promoter of the *nptII* gene is active in a wide range of taxa and has been shown to be suitable to drive fluorescent protein expression from single chromosomal insertions (Ledermann *et al.*, 2015). All bacterial strains that carried either a plasmid of the pMRE-series, Tn*5* or Tn*7*-based transposons, exhibited visible fluorescence at a single-cell resolution. This shows the application of the here constructed toolbox and its potential use to generate fluorescently-tagged bacterial communities.

The plasmids were tested in a wide range of bacterial taxa that originated from aboveground surfaces of plants (Bai *et al.*, 2015; Feil *et al.*, 2005; Innerebner *et al.*, 2011; Remus-Emsermann *et al.*, 2013a, 2016b; Schmid *et al.*, 2018; Smits *et al.*, 2010). For efficient delivery of the plasmids, conjugation using *E. coli* S17-1 was performed. The presence of the *tra* operon in this strain allows for the mobilisation of plasmids to recipient strains (Simon *et al.*, 1983), bypassing the need to produce and optimise competent cells for every recipient strain used, which can be very time and labour intensive for strains that were recently isolated. However, optimisation of conjugation conditions might be necessary to increase conjugation efficiencies.

pFru97-based pMRE plasmids could be conjugated into Pseudomonads and Enterobacteriaceae supporting previous findings (Table 4) (Miller *et al.*, 2000; Nakai *et al.*, 2001). In contrast to previous findings, the *Microbacterium* sp. Leaf320 tested in this study did not readily accept pMRE plasmids compared to other *Microbacterium* sp. tested previously (Lin *et al.*, 2012). For short term studies, or providing that the pMRE plasmids are highly stable in the used bacterial strains, they could be advantageous to other systems, since fluorescence is generally higher than after chromosomal insertion (Figure 3B, Supplementary Figure 2).

The delivery of fluorescent protein genes into bacterial isolates was most successful using the Tn*5* transposon compared to the other two delivery systems. Most members of the Proteobacteria, including members of the α- and γ-Proteobacteria, were integrating the Tn*5* transposon into their genome, which is in line with previous observations (Andersen *et al.*, 1998; Schada von Borzyskowski *et al.*, 2015). The Tn*5* transposome system, employing Tn*5* transposase bound to transposon DNA flanked by Tn*5* inverted repeats, has been shown to be suitable to construct random insertion mutations in Gram-positive bacteria (Fernandes *et al.*, 2001; Vidal *et al.*, 2009). Consequently, Tn*5* transposon integration was tested in Gram-positive bacteria as well. Since the purpose of the constructs described here is not to generate mutant libraries but to fluorescently tag bacteria, Tn*5* insertions at low frequencies would be sufficient to serve the purpose as a gene delivery tool. However, we failed to identify positive mutants in the two Gram-positive strains tested. Even though this strain selection is not exhaustive, the lack of evidence of plasmid-based Tn*5* transposon delivery into Gram-positive strains underlines the slim chances of successfully using the here proposed system in Gram-positives. In the future, additional plasmids will be developed to allow fluorescent protein gene delivery into Gram-positive bacteria. The here constructed plasmids will serve as a foundation for this work. Transposon mediated delivery tools such as Himar will be employed to deliver fluorescent protein genes to widened bacterial host range (Nilsson *et al.*, 2014).

The functionality of Tn*7* transposons is well documented in Proteobacteria. However, the here employed plasmid backbone has been predominantly used in *E. coli* and very close relatives such as *Salmonella* (McKenzie and Craig, 2006; Remus-Emsermann *et al.*, 2016a). In *E. coli*, pMRE-Tn7 plasmids are conditional suicide plasmids that fail to replicate when cells are cultivated at temperatures above 32 °C. In other bacteria, this plasmid is a suicide plasmid and is not replicating. pMRE-Tn7 plasmids carry the complete Tn*7* transposase machinery *in cis* (McKenzie and Craig, 2006). This is a great advantage compared to classical Tn*7* systems, which usually involve helper plasmids that provide the transposase machinery *in trans* (Choi and Schweizer, 2006; Lambertsen *et al.*, 2004). As the transposon machinery has to be provided on a second plasmid, the probability for both plasmids to be electroporated or conjugated into the same target cell is expected to be lower than in the system established by McKenzie and Craig (2006). Without having compared the systems of Choi and Schweizer or Lambertsen *et al.*, we expect that transposon delivery will be more successful using the here used delivery system. Even though the delivery system has previously been used in Pseudomonads (Klümper *et al.*, 2014), to our knowledge, this is the first successful application of the system as a suicide plasmid in Enterobacteriaceae and *Sphingomonas melonis* FR1 that do not replicate the plasmid backbone. To determine the successful genomic integration of the Tn*5* or Tn*7* transposons, we provide a convenient PCR-based tool that allows for screening integration events. In cases where the precise integration site needs to be determined, an arbitrary PCR should be performed as suggested by Das *et al.* (2005). To determine differences in fitness, which might be due to Tn*5* insertions into essential genes, it is advisable to perform growth experiments and competitions assays to compare wild type and mutant performances.

Using standard epifluorescence widefield microscopy, we were able to distinguish four differentially labelled bacterial strains (Figure 4). Using less common fluorescent filters and/or spectral linear unmixing approaches (Zimmermann, 2005), it should be possible to unambiguously identify five different fluorescently tagged populations. Next to *E. coli* DH5α, environmental strains also expressed fluorescent proteins to a similar degree that enabled observation of bacteria at the single-cell resolution (Figures 5 and Supplementary Figure 2).

Recently, many techniques have been developed to investigate fluorescently-tagged bacteria *in situ*, which for instance allowed the detection of 15 differentially tagged bacterial strains from dental plaques (Valm *et al.*, 2012). In combination with other experimental advances such as removing autofluorescent background from environmental samples (Peredo and Simmons, 2017; Remus-Emsermann *et al.*, 2014) and spatially explicit analysis of bacteria *in situ* (Daims *et al.*, 2006; Schmidt *et al.*, 2018), the here constructed molecular tools set the foundation to further advance *in situ* investigation of environmental bacteria.

We present a plasmid toolbox suitable to fluorescently label a broad range of bacteria for research in microbiology and microbial ecology that will be provided to the scientific community. The use of these plasmids will enable convenient tagging of many environmental bacteria and will thereby facilitate several disciplines of microbiology, such as single-cell microbiology, synthetic community, biofilm, host-microbe and microbe-microbe interactions. The ability to tag bacteria with unambiguous fluorescent colours in combination with antibiotic resistance markers to track bacterial population development at the single-cell resolution as well as on the whole population scale will be invaluable for many studies.

All plasmids constructed in this work have been made available in the convenient conjugation strain *E. coli* S17-1 that allows conjugation of the mobilisable plasmids into other bacteria at the non-profit plasmid repository Addgene (www.addgene.org).

## Acknowledgments

Raphael Ledermann and Hans-Martin Fischer (ETH Zurich) are acknowledged for contributing to the construction of pMRE-Tn5-sYFP.

## Author Contributions

RS constructed the pMRE-Tn*5* plasmid series, performed most other experiments, analysed the data and wrote the manuscript. HJ constructed the pMRE plasmid series. MB constructed the pMRE-Tn*7* plasmid series. SO performed genotyping PCRs. EB performed several mating experiments with γ-proteobacteria and contributed to other experiments. DML constructed pMRE134 and pMRE137. RD contributed materials and facilities. DR contributed to experiments, supervision, discussion, and writing. MR-E conceived and supervised the study, wrote the manuscript, contributed to some experiments and analysed data. All authors have read and approved the manuscript.

## Data availability

Plasmids and plasmid maps are available at the repository Addgene (addgene.org).

## References

- Andersen, J. B., Sternberg, C., Poulsen, L. K., Bjorn, S. P., Givskov, M., and Molin, S. (1998). New unstable variants of green fluorescent protein for studies of transient gene expression in bacteria. Appl. Environ. Microbiol. 64, 2240–2246.

- Andersson, D. I., and Hughes, D. (2011). Persistence of antibiotic resistance in bacterial populations. FEMS Microbiol. Rev. 35, 901–911.

- Bai, Y., Müller, D. B., Srinivas, G., Garrido-Oter, R., Potthoff, E., Rott, M., et al. (2015). Functional overlap of the Arabidopsis leaf and root microbiota. Nature 528, 364–369.

- Barbier, M., and Heath Damron, F. (2016). Rainbow Vectors for Broad-Range Bacterial Fluorescence Labeling. PLoS One 11, e0146827.

- Benoit, R. M., Ostermeier, C., Geiser, M., Li, J. S. Z., Widmer, H., and Auer, M. (2016). Seamless Insert-Plasmid Assembly at High Efficiency and Low Cost. PLoS One 11, e0153158.

- Bindels, D. S., Haarbosch, L., van Weeren, L., Postma, M., Wiese, K. E., Mastop, M., et al. (2017). mScarlet: a bright monomeric red fluorescent protein for cellular imaging. Nat. Methods 14, 53–56.

- Bloemberg, G. V., Wijfjes, A. H., Lamers, G. E., Stuurman, N., and Lugtenberg, B. J. (2000). Simultaneous imaging of Pseudomonas fluorescens WCS365 populations expressing three different autofluorescent proteins in the rhizosphere: new perspectives for studying microbial communities. Mol. Plant. Microbe. Interact. 13, 1170–1176.

- Carroll, D. (2011). Genome engineering with zinc-finger nucleases. Genetics 188, 773–782.

- Choi, K.-H., and Schweizer, H. P. (2006). mini-Tn7 insertion in bacteria with single attTn7 sites: example Pseudomonas aeruginosa. Nat. Protoc. 1, 153–161.

- Christen, B., Abeliuk, E., Collier, J. M., Kalogeraki, V. S., Passarelli, B., Coller, J. A., et al. (2011). The essential genome of a bacterium. Mol. Syst. Biol. 7, 528.

- Chu, J., Haynes, R. D., Corbel, S. Y., Li, P., González-González, E., Burg, J. S., et al. (2014). Non-invasive intravital imaging of cellular differentiation with a bright red-excitable fluorescent protein. Nat. Methods 11, 572–578.

- Daims, H., Lücker, S., and Wagner, M. (2006). daime, a novel image analysis program for microbial ecology and biofilm research. Environ. Microbiol. 8, 200–213.

- Das, S., Noe, J. C., Paik, S., and Kitten, T. (2005). An improved arbitrary primed PCR method for rapid characterization of transposon insertion sites. J. Microbiol. Methods 63, 89–94.

- de Lorenzo, V., Herrero, M., Jakubzik, U., and Timmis, K. N. (1990). Mini-Tn5 transposon derivatives for insertion mutagenesis, promoter probing, and chromosomal insertion of cloned DNA in gram-negative eubacteria. J. Bacteriol. 172, 6568–6572.

- Diard, M., Garcia, V., Maier, L., Remus-Emsermann, M. N. P., Regoes, R. R., Ackermann, M., et al. (2013). Stabilization of cooperative virulence by the expression of an avirulent phenotype. Nature 494, 353–356.

- Feil, H., Feil, W. S., Chain, P., Larimer, F., DiBartolo, G., Copeland, A., et al. (2005). Comparison of the complete genome sequences of Pseudomonas syringae pv. syringae B728a and pv. tomato DC3000. Proc. Natl. Acad. Sci. U. S. A. 102, 11064–11069.

- Fernandes, P. J., Powell, J. A., and Archer, J. A. (2001). Construction of Rhodococcus random mutagenesis libraries using Tn5 transposition complexes. Microbiology 147, 2529–2536.

- Gibson, D. G., Young, L., Chuang, R.-Y., Venter, J. C., Hutchison, C. A., 3rd, and Smith, H. O. (2009). Enzymatic assembly of DNA molecules up to several hundred kilobases. Nat. Methods 6, 343–345.

- Goedhart, J., von Stetten, D., Noirclerc-Savoye, M., Lelimousin, M., Joosen, L., Hink M. A., et al. (2012). Structure-guided evolution of cyan fluorescent proteins towards a quantum yield of 93%. Nat. Commun. 3, 751.

- Harder, W., Attwood, M. M., and Quayle, J. R. (1973). Methanol Assimilation by Hyphomicrobium sp. Microbiology 78, 155–163.

- Innerebner, G., Knief, C., and Vorholt, J. A. (2011). Protection of Arabidopsis thaliana against leaf-pathogenic Pseudomonas syringae by Sphingomonas strains in a controlled model system. Appl. Environ. Microbiol. 77, 3202–3210.

- Jain, A., and Srivastava, P. (2013). Broad host range plasmids. FEMS Microbiol. Lett. 348, 87–96.

- Jinek, M., Chylinski, K., Fonfara, I., Hauer, M., Doudna, J. A., and Charpentier, E. (2012). A programmable dual-RNA-guided DNA endonuclease in adaptive bacterial immunity. Science 337, 816–821.

- Klümper, U., Droumpali, A., Dechesne, A., and Smets, B. F. (2014). Novel assay to measure the plasmid mobilizing potential of mixed microbial communities. Front. Microbiol. 5, 730.

- Kremers, G.-J., Goedhart, J., van den Heuvel, D. J., Gerritsen, H. C., and Gadella, T. W. J., Jr (2007). Improved green and blue fluorescent proteins for expression in bacteria and mammalian cells. Biochemistry 46, 3775–3783.

- Kroupitski, Y., Golberg, D., Belausov, E., Pinto, R., Swartzberg, D., Granot, D., et al. (2009). Internalization of Salmonella enterica in leaves is induced by light and involves chemotaxis and penetration through open stomata. Appl. Environ. Microbiol. 75, 6076–6086.

- Lagendijk, E. L., Validov, S., Lamers, G. E. M., de Weert, S., and Bloemberg, G. V. (2010). Genetic tools for tagging Gram-negative bacteria with mCherry for visualization in vitro and in natural habitats, biofilm and pathogenicity studies. FEMS Microbiol. Lett. 305, 81–90.

- Lambertsen, L., Sternberg, C., and Molin, S. (2004). Mini-Tn7 transposons for site-specific tagging of bacteria with fluorescent proteins. Environ. Microbiol. 6, 726–732.

- Lambert, T. (2018). Tlambert03/Fpbase: V1.1.0. Zenodo doi:10.5281/ZENODO.1244328.

- Lau, B. T. C., Malkus, P., and Paulsson, J. (2013). New quantitative methods for measuring plasmid loss rates reveal unexpected stability. Plasmid 70, 353–361.

- Ledermann, R., Bartsch, I., Remus-Emsermann, M. N., Vorholt, J. A., and Fischer, H.-M. (2015). Stable Fluorescent and Enzymatic Tagging of Bradyrhizobium diazoefficiens to Analyze Host-Plant Infection and Colonization. Mol. Plant. Microbe. Interact. 28, 959–967.

- Lin, L., Guo, W., Xing, Y., Zhang, X., Li, Z., Hu, C., et al. (2012). The actinobacterium Microbacterium sp. 16SH accepts pBBR1-based pPROBE vectors, forms biofilms, invades roots, and fixes N2 associated with micropropagated sugarcane plants. Appl. Microbiol. Biotechnol. 93, 1185–1195.

- Liu, H., Bouillaut, L., Sonenshein, A. L., and Melville, S. B. (2013). Use of a mariner-based transposon mutagenesis system to isolate Clostridium perfringens mutants deficient in gliding motility. J. Bacteriol. 195, 629–636.

- McKenzie, G. J., and Craig, N. L. (2006). Fast, easy and efficient: site-specific insertion of transgenes into enterobacterial chromosomes using Tn7 without need for selection of the insertion event. BMC Microbiol. 6, 39.

- Miller, W. G., Leveau, J. H., and Lindow, S. E. (2000). Improved gfp and inaZ broad-host-range promoter-probe vectors. Mol. Plant. Microbe. Interact. 13, 1243–1250.

- Million-Weaver, S., Alexander, D. L., Allen, J. M., and Camps, M. (2012). Quantifying plasmid copy number to investigate plasmid dosage effects associated with directed protein evolution. Methods Mol. Biol. 834, 33–48.

- Monier, J.-M., and Lindow, S. E. (2005). Spatial organization of dual-species bacterial aggregates on leaf surfaces. Appl. Environ. Microbiol. 71, 5484–5493.

- Nakai, J., Ohkura, M., and Imoto, K. (2001). A high signal-to-noise Ca(2+) probe composed of a single green fluorescent protein. Nat. Biotechnol. 19, 137–141.

- Nilsson, M., Christiansen, N., Høiby, N., Twetman, S., Givskov, M., and Tolker-Nielsen, T. (2014). A mariner transposon vector adapted for mutagenesis in oral streptococci. Microbiologyopen 3, 333–340.

- Parks, A. R., and Peters, J. E. (2007). Transposon Tn7 is widespread in diverse bacteria and forms genomic islands. J. Bacteriol. 189, 2170–2173.

- Peredo, E. L., and Simmons, S. L. (2017). Leaf-FISH: Microscale Imaging of Bacterial Taxa on Phyllosphere. Front. Microbiol. 8, 2669.

- Peters, J. E. (2014). Tn7. Mirobiol Spectr 2. doi:10.1128/microbiolspec.MDNA3-0010-2014.

- Ramirez-Mata, A., Pacheco, M. R., Moreno, S. J., Xiqui-Vazquez, M. L., and Baca, B.E. (2018). Versatile use of Azospirillum brasilense strains tagged with egfp and mCherry genes for the visualization of biofilms associated with wheat roots. Microbiol. Res. 215, 155–163.

- Remus-Emsermann, M. N. P., Gisler, P., and Drissner, D. (2016a). MiniTn7-transposon delivery vectors for inducible or constitutive fluorescent protein expression in Enterobacteriaceae. FEMS Microbiol. Lett. 363. doi:10.1093/femsle/fnw178.

- Remus-Emsermann, M. N. P., Kim, E. B., Marco, M. L., Tecon, R., and Leveau, J. H. J. (2013a). Draft Genome Sequence of the Phyllosphere Model Bacterium Pantoea agglomerans 299R. Genome Announc. 1. doi:10.1128/genomeA.00036-13.

- Remus-Emsermann, M. N. P., Kowalchuk, G. A., and Leveau, J. H. J. (2013b). Single-cell versus population-level reproductive success of bacterial immigrants to pre-colonized leaf surfaces. Environ. Microbiol. Rep. 5, 387–392.

- Remus-Emsermann, M. N. P., Lücker, S., Müller, D. B., Potthoff, E., Daims, H., and Vorholt, J. A. (2014). Spatial distribution analyses of natural phyllosphere-colonizing bacteria on Arabidopsis thaliana revealed by fluorescence in situ hybridization. Environ. Microbiol. 16, 2329–2340.

- Remus-Emsermann, M. N. P., and Schlechter, R. O. (2018). Phyllosphere microbiology: at the interface between microbial individuals and the plant host. New Phytol. 218, 1327–1333.

- Remus-Emsermann, M. N. P., Schmid, M., Gekenidis, M.-T., Pelludat, C., Frey, J. E., Ahrens, C. H., et al. (2016b). Complete genome sequence of Pseudomonas citronellolis P3B5, a candidate for microbial phyllo-remediation of hydrocarbon-contaminated sites. Stand. Genomic Sci. 11, 75.

- Reznikoff, W. S. (2008). Transposon Tn5. Annu. Rev. Genet. 42, 269–286.

- Rodriguez, M. D., Paul, Z., Wood, C. E., Rice, K. C., and Triplett, E. W. (2017). Construction of Stable Fluorescent Reporter Plasmids for Use inStaphylococcus aureus. Front. Microbiol. 8, 2491.

- Sambrook, J., Fritsch, E. F., Maniatis, T., and Others (1989). Molecular cloning: a laboratory manual. Cold spring harbor laboratory press.

- Schada von Borzyskowski, L., Remus-Emsermann, M., Weishaupt, R., Vorholt, J. A., and Erb, T. J. (2015). A set of versatile brick vectors and promoters for the assembly, expression, and integration of synthetic operons in Methylobacterium extorquens AM1 and other alphaproteobacteria. ACS Synth. Biol. 4, 430–443.

- Schindelin, J., Arganda-Carreras, I., Frise, E., Kaynig, V., Longair, M., Pietzsch, T., et al. (2012). Fiji: an open-source platform for biological-image analysis. Nat. Methods 9, 676–682.

- Schmid, M., Frei, D., Patrignani, A., Schlapbach, R., Frey, J. E., Remus-Emsermann, M. N. P., et al. (2018). Pushing the limits of de novo genome assembly for complex prokaryotic genomes harboring very long, near identical repeats. bioRxiv, 300186.

- Schmidt, H., Nunan, N., Höck, A., Eickhorst, T., Kaiser, C., Woebken, D., et al. (2018). Recognizing Patterns: Spatial Analysis of Observed Microbial Colonization on Root Surfaces. Front. Environ. Sci. Eng. China 6. doi:10.3389/fenvs.2018.00061.

- Simon, R., Priefer, U., and Pühler, A. (1983). A Broad Host Range Mobilization System for In Vivo Genetic Engineering: Transposon Mutagenesis in Gram Negative Bacteria. Biotechnology 1, 784.

- Smith, M. A., and Bidochka, M. J. (1998). Bacterial fitness and plasmid loss: the importance of culture conditions and plasmid size. Can. J. Microbiol. 44, 351–355.

- Smits, T. H. M., Rezzonico, F., Kamber, T., Blom, J., Goesmann, A., Frey, J. E., et al. (2010). Complete genome sequence of the fire blight pathogen Erwinia amylovora CFBP 1430 and comparison to other Erwinia spp. Mol. Plant. Microbe. Interact. 23, 384–393.

- Stewart, P. S., and Costerton, J. W. (2001). Antibiotic resistance of bacteria in biofilms. Lancet 358, 135–138.

- Suarez, A., Güttler, A., Strätz, M., Staendner, L. H., Timmis, K. N., and Guzmán, C. A. (1997). Green fluorescent protein-based reporter systems for genetic analysis of bacteria including monocopy applications. Gene 196, 69–74.

- Subach, O. M., Cranfill, P. J., Davidson, M. W., and Verkhusha, V. V. (2011). An enhanced monomeric blue fluorescent protein with the high chemical stability of the chromophore. PLoS One 6, e28674.

- Summers, D. K. (1991). The kinetics of plasmid loss. Trends Biotechnol. 9, 273–278.

- Tecon, R., and Leveau, J. H. J. (2012). The mechanics of bacterial cluster formation on plant leaf surfaces as revealed by bioreporter technology. Environ. Microbiol. 14, 1325–1332.

- Thomas, C. M., and Nielsen, K. M. (2005). Mechanisms of, and barriers to, horizontal gene transfer between bacteria. Nat. Rev. Microbiol. 3, 711–721.

- Tolker-Nielsen, T., and Molin, S. (2000). Spatial Organization of Microbial Biofilm Communities. Microb. Ecol. 40, 75–84.

- Valm, A. M., Mark Welch, J. L., and Borisy, G. G. (2012). CLASI-FISH: principles of combinatorial labeling and spectral imaging. Syst. Appl. Microbiol. 35, 496–502.

- Vidal, J. E., Chen, J., Li, J., and McClane, B. A. (2009). Use of an EZ-Tn5-based random mutagenesis system to identify a novel toxin regulatory locus in Clostridium perfringens strain PLoS One 4, e6232.

- Whitaker, W. R., Shepherd, E. S., and Sonnenburg, J. L. (2017). Tunable Expression Tools Enable Single-Cell Strain Distinction in the Gut Microbiome. Cell 169, 538–546.e12.

- Zengerer, V., Schmid, M., Bieri, M., Müller, D. C., Remus-Emsermann, M. N. P., Ahrens, C. H., et al. (2018). Pseudomonas orientalis F9: A Potent Antagonist against Phytopathogens with Phytotoxic Effect in the Apple Flower. Front. Microbiol. 9, 145.

- Zimmermann, T. (2005). “Spectral Imaging and Linear Unmixing in Light Microscopy,” in Microscopy Techniques Advances in Biochemical Engineering/Biotechnology., ed. J. Rietdorf (Berlin, Heidelberg: Springer Berlin Heidelberg), 245–265.

